# *DRP1* depletion protects NK cells against hypoxia-induced dysfunction

**DOI:** 10.1101/2025.06.23.661011

**Authors:** Tias Verhezen, Astrid Van den Eynde, Peter Verstraelen, Laura Gehrcken, Gabriele Palmiotto, Ho Wa Lau, Winnok de Vos, Sanne van der Heijden, Louize Brants, Jöran Melis, Jonas Van Audenaerde, Steven Van Laere, Filip Lardon, Christophe Deben, An Wouters, Evelien Smits, Jorrit De Waele

## Abstract

Hypoxia within the tumor microenvironment poses a major barrier to the efficacy of NK cell-based immunotherapies for solid tumors. In this study, we investigated the influence of hypoxia on NK cell function and mitochondria. We found that hypoxia reduced NK cell cytotoxicity, mitochondrial content, and membrane potential, while increasing mtROS and inducing broad transcriptional changes in metabolic and stress response pathways. CAR engineering with CD70 and IL-15, while designed to enhance persistence and metabolic fitness, did not prevent hypoxia-induced impairment. Given the mitochondrial disruption, we then explored whether DRP1 ablation could mitigate hypoxia-induced dysfunction. Pharmacological inhibition of DRP1 restored mitochondrial content and cytotoxic function. To confirm the role of DRP1, we generated CRISPR-Cas9-mediated DRP1 KO NK cells, which preserved mitochondrial load and membrane potential under hypoxia. When armed with CD70-CAR-IL-15, DRP1^KO^ cells retained cytotoxic activity under hypoxic conditions. These findings show that DRP1 inactivation can support NK cell function in hypoxic environments, and that metabolic engineering may enhance CAR NK cell efficacy in solid tumors.

**Graphical abstract:** NK cells become dysfunctional in hypoxic conditions, while DRP1^KO^ NK cells retain their function.

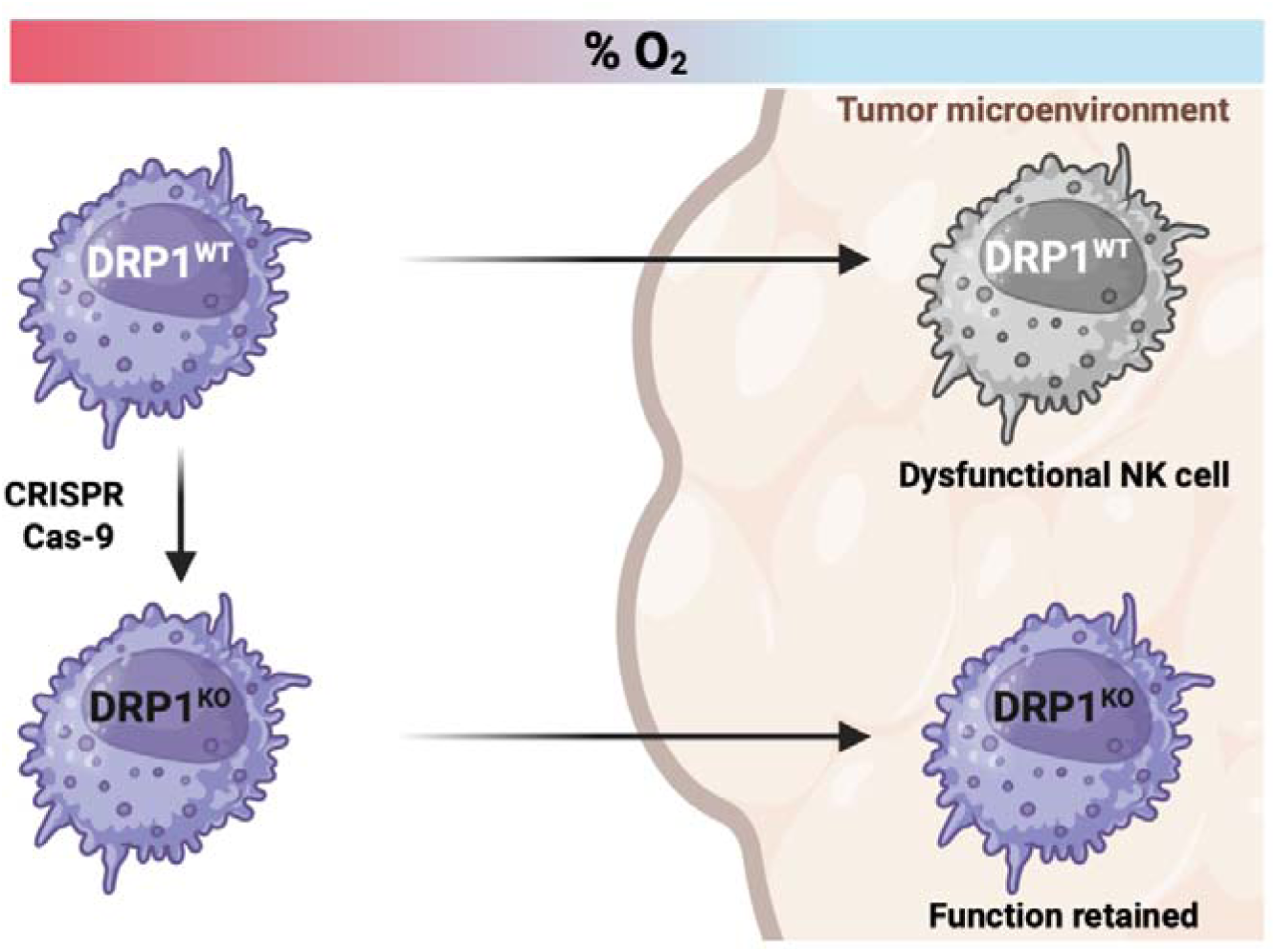

## Introduction

Cell and gene therapies are revolutionizing the way cancer is treated. CAR T cell therapies have demonstrated the ability to induce durable remissions or even achieve cures in patients with blood cancers that were in final stages, including leukemia and lymphoma^1,2^. Despite these hopeful results in hematological malignancies, the application of CAR therapies has been more challenging in solid tumors^3,4^. Up to 90 percent of adult cancers consist of solid tumors, presenting a big need for new therapies^5^. Solid tumors have a complex tumor microenvironment (TME) that creates barriers for immune cell infiltration and function. This immune-suppressive niche is low in nutrients, highly acidic and hypoxic^6,7^. Accumulating data indicates that T cells and NK cells entering this hostile environment are rendered dysfunctional^8–12^.

NK cells are cytotoxic cells from the innate immune system that can recognize and eradicate tumor cells swiftly and without prior sentitization^13^. Their distinct advantages over T cells, including a superior safety profile and potential for allogeneic therapy, make NK cell treatments more economically and logistically feasible, broadening patient accessibility^14^. In addition, NK cells can eradicate tumor cells beyond their CAR construct due to their innate properties, being a major benefit in the heterogeneous TME of solid tumors. The first-in-human clinical trial with CAR NK cells has shown promising results in CD19^+^ hematological malignancy^15^. Until now, efforts to enhance NK cell efficacy in solid cancer therapy have mainly focused on blocking inhibitory receptors, CAR engineering, NK cell engagers or cytokine stimulation. However, recognizing the profound impact of the TME on immunometabolism, recent approaches aim to combine immunotherapy with metabolic manipulation to bolster cellular function^16^.

Hypoxia is a defining feature of most solid tumor types, arising from the rapid proliferation and altered metabolism of cancer cells, coupled with poorly organized and dysfunctional vasculature^6,17^. This imbalance between oxygen supply and demand creates steep oxygen gradients, with partial pressures dropping below 10 mmHg^18^. In normal tissues, the oxygen tension is in the range of 24-66LmmHg^19^. Hypoxia profoundly influences the behavior of cancer cells, leading to tumor progression, and contributes to resistance against chemotherapy, radiotherapy, and immunotherapy through a variety of mechanisms^17^. Clinically, hypoxia is associated with poor prognosis across a wide range of solid tumors^17^. Importantly, hypoxia also poses a major challenge for the efficacy of immune-based therapies, including adoptive cell therapies such as those using NK cells. The hypoxic TME can impair NK cell metabolism, survival, and effector functions, limiting their ability to eliminate tumor cells^8,12,20–22^. Understanding how hypoxia shapes NK cell biology is therefore critical for optimizing the design and application of NK cell-based immunotherapies in solid cancers.

In this study, we investigated the influence of hypoxia on NK cells, including in the context of cytokine-armored fourth generation CAR NK cells. We confirmed that hypoxia impairs mitochondrial function and cytotoxicity in NK cells. This dysfunction was also observed in CAR NK cells, despite the incorporation of IL-15 to enhance persistence and metabolic fitness. Building on the known role of DRP1 in hypoxia-induced mitochondrial fragmentation, and the importance of mitochondrial fitness in NK cell cytotoxicity, we hypothesized that genetic ablation of DRP1 could preserve NK cell function under metabolic stress^8^. To test this, we investigated both pharmacological inhibition and CRISPR-Cas9-mediated knockout (KO) of DRP1 in NK cells exposed to hypoxia. Indeed, DRP1^KO^ NK cells maintained mitochondrial content and cytotoxic potential in hypoxic conditions.

## Materials and Methods

### Cell lines and culture conditions

The NK-92 and Raji cell lines were purchased from the German Collection of Microorganisms and Cell Cultures. The PANC-1 cell line was purchased from the American Type Culture Collection. The HeLa and K562 cell lines were kindly provided by Dr. Eva González Suárez (Spanish National Cancer Research Centre) and Dr. Eva Lion (University of Antwerp), respectively.

NK-92 cells were cultured in GlutaMAX alpha Minimum Essential Medium (α-MEM; Life Technologies) supplemented with 12.5% Fetal Bovine Serum (FBS; Life Technologies), 12.5% horse serum (Life Technologies), 2mM L-glutamine (Life Technologies), 1% Penicillin/Streptomycin (P/S; Life Technologies) and 150 U/mL recombinant IL-2 (ImmunoTools), as described before^23^. Raji cells were cultured in Roswell Park Memorial Institute Medium (Life Technologies) supplemented with 10% FBS, 2mM L-glutamine, and 1% P/S. Panc-1 and HeLa cells were cultivated in Dulbecco Modified Eagle Medium (DMEM; Life Technologies) supplemented with 10% FBS, 2mM L-glutamine, and 1% P/S. Cell cultures were maintained in exponential growth at 37 °C in a humidified incubator with 5% COL. Regular testing using the MycoAlert Mycoplasma Detection Kit (Lonza) confirmed the absence of Mycoplasma contamination, and cell identity was validated through short tandem repeat profiling.

### Hypoxic environment

In this study, we used 1% O_2_ to model tumor hypoxia *in vitro*. This level is commonly used in cancer research to mimic the hypoxic microenvironment of solid tumors, where oxygen tensions often fall below 10 mmHg (∼1-1.5% O_2_)^12,18^. Furthermore, 1% O_2_ is sufficient to activate hypoxia-inducible pathways such as HIF-1α stabilization, providing a reliable model for studying hypoxic cellular responses^24^. Hypoxic conditions were reached by placing DRP1^WT^ and DRP1^KO^ NK-92 cells in H45 HEPA Hypoxystations (Don Whitley Scientific) at 1% O_2_ and 5% CO_2_ at 37 °C. Unless stated differently, cells were kept in hypoxia for 48 h before downstream experiments.

### Growth curve NK-92 cells

NK-92 cell expansion was determined by calculating the fold increase in cell concentration. NK cells were initially seeded at a concentration of 0.333 × 10L cells/mL in culture medium. After an incubation period of 3-4 days, the final cell concentration was measured using a Bio-Rad automated cell counter with Trypan Blue viability staining. This was repeated over the entire culture period.

### TCGA analysis

Gene expression data (RSEM format) for 32 different cancer types vouching for 10,071 individual cancer samples that are part of the pan-cancer analysis of whole genomes (PCAWG) series were downloaded from the cBioPortal for cancer genomics (https://www.cbioportal.org). Expression data were filtered to include genes with raw counts above 10 in at least 10% of the cases per cancer type, followed by library size scaling (count per million reads), log2 transformation and quantile normalization. Hypoxia scores were generated using gene set variation analysis (GSVA) based on a set of 15 hypoxia-related genes (VEGFA, PGAM1, ENO1, LDHA, TPI1, P4HA1, MRPS17, ADM, NDRG1, TUBB6, ALDOA, MIF, SLC2A1, CDKN3, ACOT7) identified by Buffa *et al.* (2010) in a meta-analysis across different cancer types^25^. Resulting hypoxia scores were compared between different tumor types.

### Generation of DRP1^KO^ NK cells via CRISPR-Cas9

DRP1^KO^ NK cells were generated with CRISPR-Cas9 editing and validated by PCR amplification and flow cytometry staining as described previously^26^. Briefly, 1 × 10^6^ IL-2 pre-activated NK cells were nucleofected with CRISPR-Cas9 multi-guide ribonucleoprotein (RNP) complexes. The following sgRNA (synthetic guide RNA) sequences were used to target the *DRP1* gene: (a) AUAUUCUGUUUUCAGAGCAG; (b) UUCCAGUACCUCUGGGAAGC; (c) GAAGAUAAACGGAAAACAAC. After 48 h, cells were harvested for genetic validation using PCR amplification. The following PCR primers were used amplify the targeted *DRP1* sequence: forward, AGTCTCTGCACTAATTTTTCCTTT; and reverse, TGCATACTACTTCTCACAGGACTT.

### Flow cytometry

*Co-culture with PKH67-stained K562 and Raji cells* - Cytotoxic activity towards the K562 and Raji cell lines was assessed by co-culturing NK cells with K562 or Raji cells in a 5:1 effector:target ratio in sterile FACS tubes (VWR). Prior to start, tumor cells were labeled using PKH67 (Sigma-Aldrich) according to the manufacturer’s instructions. After 4 h, co-cultures were stained with 7-AAD and Annexin V-PE (BD Biosciences). Tumor cell survival was measured on a CytoFLEX flow cytometer (Beckman Coulter). Fig. **S1C** and **S2B** show the gating strategy used to distinguish between K562/Raji and NK cells, and live, apoptotic or dead K562/Raji cells.

*TMRM and MitoSOX staining* - Mitochondrial membrane potential of NK cells was quantified by Tetramethylrhodamine, Methyl Ester, Perchlorate (TMRM) staining and mtROS levels were visualized by MitoSOX mitochondrial superoxide indicators (both ThermoFisher Scientific). NK cells were harvested in U-bottom 96-well plates and resolved in serum-free alpha-MEM medium. NK cells were incubated with 100 nM TMRM or 1 µM MitoSOX for 30 min at 37 °C, washed with FACS buffer, and stained with with LIVE/DEAD™ Fixable Near-IR Dead Cell Stain (Thermo Fisher Scientific) for 15 min. TMRM and MitoSOX signals were measured using a NovoCyte Quanteon Flow Cytometer, Mean Fluorescence Intensity (MFI) was used to check TMRM and MitoSOX signals.

*DRP1 staining* - NK cells were harvested and washed with PBS-EDTA before staining with LIVE/DEAD™ Fixable Near-IR Dead Cell Stain (Thermo Fisher Scientific) for 15 min. Following incubation, cells were washed, fixed and permeabilized using eBioscience Foxp3 fixation and permeabilization buffer kit according to the instructions of the manufacturer (Thermo Fisher Scientific). To block nonspecific binding, an intracellular blocking step was included with blocking buffer (permeabilization buffer with 10% human serum). Cells were then stained with Phospho-DRP1 (Ser616) Antibody (source: rabbit) (Cell Signaling Technology, Cat. No. 3455) in blocking buffer for 1 h, at 4 °C. This was followed by a washing step and a 30 min incubation with 1/500 Goat anti-Rabbit IgG secondary antibody (Alexa Fluor™ 647) (Invitrogen) in permeabilization buffer. DRP1 signal was measured using a NovoCyte Quanteon Flow Cytometer, Mean Fluorescence Intensity was used to check DRP1 expression.

*CD70 expression on cancer cells and CD70-targeting CAR expression on NK cells* - Surface expression of the CD70-CAR-IL-15 constructs on NK cells was evaluated via flow cytometry by staining with a CD27-PE monoclonal antibody (Clone O323, Cell Signaling Technology) for 15 min. Expression of CD70 on tumor and CAF cell lines was determined by staining with a CD70-PE mAb (Clone J606, BD Biosciences) for 15 min. To exclude non-viable cells, the mAB stainings were combined with the LIVE/DEAD™ Fixable Near-IR Dead Cell Stain (Thermo Fisher Scientific). CD27 and CD70 expression were measured on a NovoCyte Quanteon Flow Cytometer.

### RNA sequencing

For RNA sequencing (RNAseq), DRP1^WT^ and DRP1^KO^ NK cells were harvested after 24 h of culturing in hypoxia or normoxia, washed with cold PBS, and stored as dry pellets at −80 °C. NK cells were subjected to 24 h of hypoxia to ensure preservation of cell viability and transcriptional activity, which could be too low after 48 h of hypoxic culturing. Afterwards, RNA was extracted using RNeasy midi kit (Qiagen). For removal of gDNA, RNAse-free DNAse treatment was performed. RNA concentration and purity were checked using the Qubit RNA BR Assay Kit on Qubit 4 Fluorometer (ThermoFisher) and NanoDrop ND-1000 (ThermoFisher), respectively. Samples were frozen at −80 °C and delivered to the Genomics Core Leuven for transcriptome sequencing using Lexogen QuantSeq 3’ FWD library preparation kit for Illumina on a Hiseq400 SR50 line with a minimum of 2 M reads per sample. Downstream analysis and plotting were performed using the Omics Playground tool (Big Omics Analytics) and RStudio.

### SCENITH assay

The SCENITH assay (SCENITH reagents kit; https://www.scenith.com) was performed as previously described by Argüello *et al.^27^*. Briefly, 0.25 × 10^6^ DRP1^WT^ or DRP1^KO^ NK cells were plated in a U-bottom, 96-well plate, treated with control (DMSO), 2-Deoxy-D-Glucose (2-DG; 100 mM), oligomycin (Oligo; 1 μM), or a combination of the inhibitors at the same final concentrations, for 30 minutes at 37 °C. Following treatment with metabolic inhibitors, puromycin (Puro; 10 μg/mL) was added during the last 30 minutes. Cells were then washed with cold PBS and stained with LIVE/DEAD™ Fixable Aqua Dead Cell Stain Kit (Thermo Fisher Scientific) for 15 minutes. Following incubation, cells were washed, fixed and permeabilized using eBioscience Foxp3 fixation and permeabilization buffer kit (Thermo Fisher Scientific). Intracellular block was performed with human serum (10%) and intracellular staining of puromycin with an anti-puromycin antibody (1:50) was then performed for 1 hour in permeabilization buffer with 10% human serum at 4 °C. Puromycin signal was measured using a NovoCyte Quanteon Flow Cytometer, MFI was used to calculate the SCENITH parameters.

Calculations used for SCENITH parameters include the following (with the puromycin MFI of each condition):

a. Glucose dependence = 100 (Control – 2DG)/(Control – [2-DG + Oligo]);
b. mitochondrial dependence = 100 (Control – Oligo)/(Control – [2-DG + Oligo]); and
c. glycolytic capacity = 100 − mitochondrial dependence.

### Confocal microscopy

*Sample preparation* - Following incubation under normoxic or hypoxic conditions, DRP1^KO^ and DRP1^WT^ NK cells were harvested, washed, and fixed in 2% PFA for 10 minutes. Subsequently, 1 × 10^5^ cells (at a concentration of 1 × 10^6^ cells/mL) were mounted onto glass slides using a Cytospin device (ThermoFisher) at 1500 rpm for 5 min to ensure uniform cell distribution. To contain the sample area, a hydrophobic barrier was drawn around the mounted cells using a ReadyProbes™ Hydrophobic Barrier Pap Pen. Cells were permeabilized with blocking buffer (PBS supplemented with 0.05% Thimerosal (Fluka), 0.01% NaN3 (Acros), 0.1% Bovine serum albumin (Sigma) and 10% Horse Serum (Gibco)) containing 1% Triton X-100 (Sigma) for 5 min. For mitochondrial staining, cells were incubated overnight with a primary anti-TOMM20 antibody (Mouse monoclonal, BD 612278) diluted 1:250 in blocking buffer (4 °C). The following day, samples were washed with PBS and incubated with a Goat-anti-Mouse-AlexaFluor488-conjugated secondary antibody (1:500 dilution) along with Cellmask (1:5000 dilution) in blocking buffer for 2 h at room temperature. To visualize cell nuclei, cells were incubated with DAPI (Sigma D9542; 5 µg/ml in PBS) for 10 min at room temperature and washed with PBS.

*Image quantification* - High-resolution confocal Z-stacks (spacing: 0.1 µm) were acquired on a Nikon CSU-W1-SoRa spinning disk confocal with a 100X silicone immersion objective (NA 1.35), using standard filters for blue (DAPI), green (AlexaFluor488) and far red (CellMask). Three-dimensional image analysis was carried out in Arivis software (Zeiss). Briefly, the DAPI channel was segmented with a blob finder to delineate nuclei, which were then used as seeds to segment the cell based on the CellMask channel. Within the cytoplasm, i.e., cell minus nucleus, mitochondria were detected in the TOMM20 channel. Mitochondrial occupancy was calculated as the volume of mitochondria divided by the volume of the cytoplasm. Numerical data was plotted and analyzed in RStudio. One data point represents one cell, originating from 4 independent experiments.

### Generation of CD70-targeting IL15-expressing CAR NK cells

CD70-CAR-IL-15 constructs were designed and created as previously described^23^. *In vitro* transcription of CD70-CAR-IL-15-encoding messenger RNA (mRNA) was performed using the mMESSAGE mMACHINE T7 transcription kit (Life Technologies) following manufacturer’s instructions. DRP1^WT^ or DRP1^KO^ NK-92 cells were electroporated using a 4D Nucleofector device (Lonza) in the presence of 4 μg CD70-CAR-IL-15-encoding mRNA per 1 million DRP1^WT^ or DRP1^KO^ NK cells.

### xCelligence coculture

Longitudinal cytotoxic activity of CD70-CAR-IL-15 DRP1^WT^ and DRP1^KO^ NK cells towards CD70+ cell lines (HeLa, Panc-1 and Raji) was analyzed using the xCELLigence Real-Time Cell Analysis (RTCA; Agilent) that records cell viability and growth by impedance measurements. Seeding density was optimized for each cell line to ensure continuous growth until the end of the assay. Target cells were seeded in gold-coated 16-well plates and background impedance of the plates was measured before seeding of the target cells. After a 24 h incubation to allow proper adhesion to the plates, cell lines were treated with CD70-CAR-IL-15 DRP1^WT^ and DRP1^KO^ NK cells in a 5:1 effector:target ratio (based on the amount of target cells seeded at day 0), or left untreated. The impedance signal was monitored by automated measurements every 15 min starting from cell seeding and ending 24 h after treatment. Measurement of the impedance was expressed as Cell Index and normalized to 1 after starting the co-culture.

### Statistics

Statistical analyses were performed using GraphPad Prism v10.2.2, JMP Pro v17.0.0 (SAS Institute Inc.), and RStudio. To compare two groups, either paired t-tests or Wilcoxon signed-rank tests were used, depending on data distribution and sample size. For comparisons involving more than two groups, one-way ANOVA or Linear Mixed Models were applied. Tukey’s post hoc correction was used for all pairwise comparisons to adjust for multiple testing. A p-value < 0.05 was considered statistically significant.

## Results

### Hypoxia impairs NK cell cytotoxicity and is associated with mitochondrial dysfunction

Improving NK-cell performance in the hypoxic TME requires a clear understanding of how low oxygen alters their metabolism and cytotoxicity. We therefore modeled TME-like oxygen levels by culturing NK-92 cells at 1% OL for 48 hours, with cells maintained at 21% OL serving as normoxic controls. As shown in **Fig. 1A**, hypoxia significantly impaired NK cell cytotoxicity against K562 target cells, confirming that low oxygen compromises effector function.

**Figure 1.**
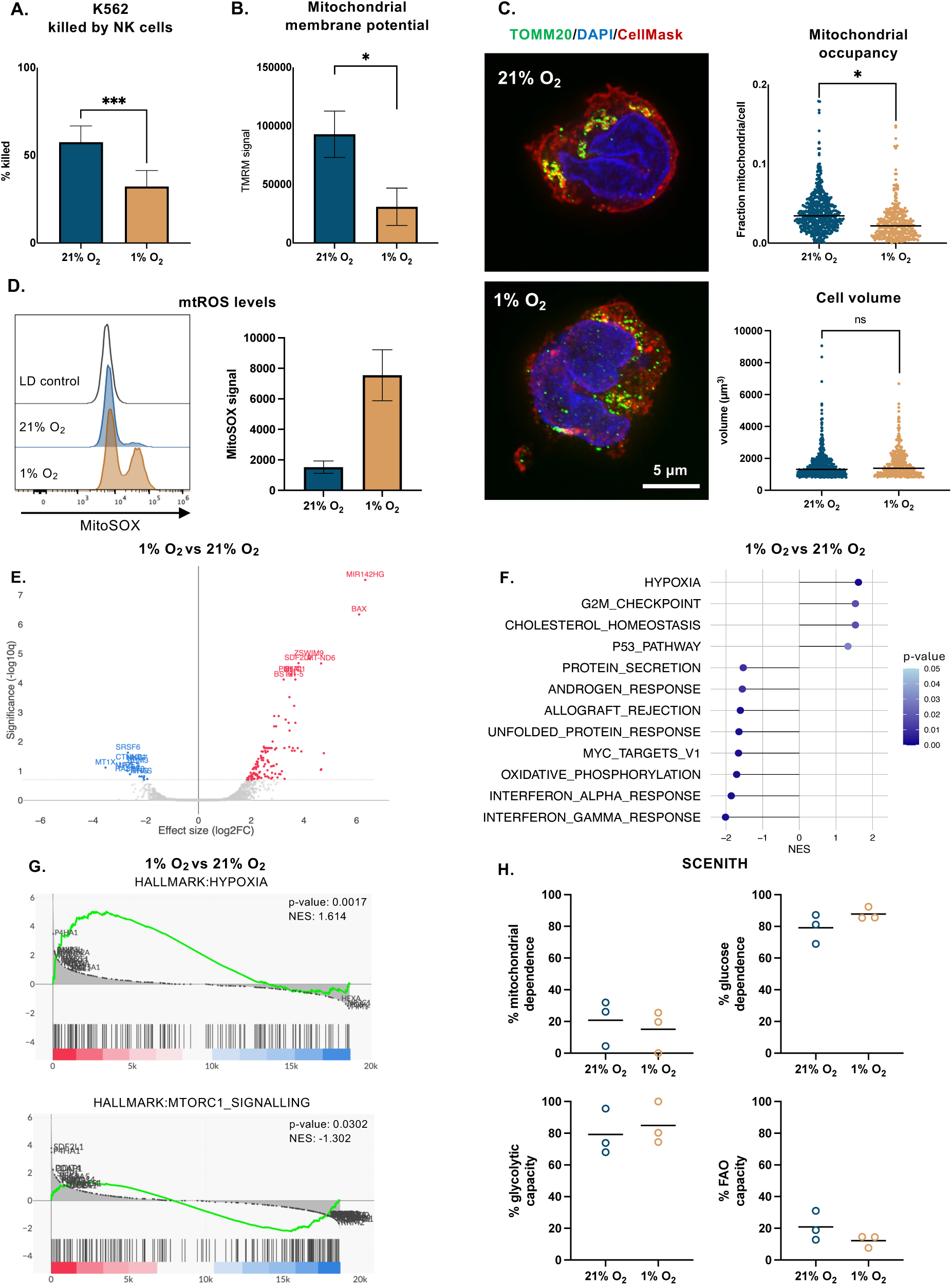
Hypoxia affects NK cell killing potential, gene expression, and metabolism. A. Cytotoxic activity (% K562 cells killed) of NK cells cultured in normoxia (21% O₂) or hypoxia (1% O₂), measured by flow cytometry; B. Mitochondrial membrane potential measured by TMRM staining in NK cells after 24 h in normoxia or hypoxia, analyzed by flow cytometry; C. Representative confocal microscopy images of NK cells showing mitochondria (TOMM20), nucleus (DAPI), and cytoplasm (CellMask) after 24 h in normoxic or hypoxic conditions, quantification of cell volume and mitochondrial occupancy is based on the images in Fig. S1B, each point represents one cell; D. mtROS levels after normoxic/hypoxic culturing as determined by mitoSOX staining; E. Volcano plot of differentially expressed genes identified between the hypoxia treated and normoxia control cells; F. Hallmark gene set enrichment analysis (GSEA) showing the top up- and downregulated pathways in hypoxia, visualized as a lollipop plot; G. GSEA enrichment plots showing increased expression of hypoxia-related genes and reduced mTORC1 signaling in hypoxic NK cells; H. SCENITH-based analysis of NK cell metabolic activity under normoxia and hypoxia, showing corresponding metabolic phenotypes. *p < 0.05; ***p < 0.001; ns = not significant. FAO: Fatty Acid Oxidation.

To identify potential metabolic contributors to this dysfunction, we next evaluated mitochondrial structure and activity in NK cells exposed to hypoxia. Mitochondrial occupancy was not affected after 24 hours, but a significant reduction was observed at 48 hours, justifying our use of this time point in subsequent experiments (**Fig. S1A, B**). Under these conditions, mitochondrial occupancy was markedly reduced despite no change in cell volume (**Fig. 1C and S1B** for representative images). Moreover, mitochondrial membrane potential, a proxy for mitochondrial activity, was significantly lower in NK cells cultured under hypoxia **(Fig. 1B),** and mitochondrial ROS (mtROS) levels were strongly elevated **(Fig. 1D)**.

To further understand how hypoxia alters NK cell metabolism, we performed RNA sequencing on NK cells cultured under normoxic and hypoxic conditions. Differential gene expression analysis revealed a distinct transcriptional profile in hypoxic NK cells, with upregulation of genes involved in stress and metabolic adaptation **(Fig. 1E)**. Among the most significantly upregulated genes in hypoxia, we identified MIR142HG, a non-coding RNA molecule implicated in NK cell immune regulation and metabolic adaptation^28^. Additionally, the pro-apoptotic gene BAX was significantly upregulated, suggesting potential stress-induced apoptotic priming. In **Fig. 1F**, the top enriched gene sets/hallmarks are shown. As expected, hypoxia-exposed NK cells exhibited significant enrichment of hypoxia hallmark genes. Conversely, genes associated with mTORC1 signaling, interferon responses, and OXPHOS were significantly downregulated (**Fig. 1F, G**). mTORC1 signaling plays a central role in regulating NK cell metabolism and effector function, while interferon responses are essential for immune surveillance and cytotoxicity^29^. The downregulation of OXPHOS-related genes is consistent with the reduced mitochondrial load and membrane potential observed in hypoxic NK cells (**Fig. 1B, C, F**).

To functionally validate these shifts in metabolic programming, we performed SCENITH, a flow cytometry-based assay that quantifies protein synthesis in response to metabolic pathway inhibition^27^. Hypoxia caused an approximately 4.5-fold reduction in puromycin MFI in control-treated cells, indicating a marked suppression of global protein translation, which is expected in a hypoxic environment^30^ (**Fig. S1D**). In addition, glycolytic capacity and glucose dependency were increased in hypoxia, while mitochondrial dependence was decreased, which is consistent with a reduced reliance on OXPHOS, as indicated by the RNA-seq data (**Fig. 1F, H**). This reflects the expected glycolytic adaptation in hypoxia at the expense of mitochondrial respiration^31^.

Together, these observations indicate that hypoxia places significant metabolic and survival stress on NK cells, predominantly through mitochondrial dysfunction, leading to functional impairment.

### CD70-CAR-IL-15-engineering is insufficient to overcome hypoxic dysfunction

While NK cells possess intrinsic antitumor activity, cytokine-armored CAR engineering is often required to maximize their anti-cancer efficacy by enhancing target specificity and persistence^32^. We previously showed that co-expression of IL-15 with a CD70-directed CAR enables robust anti-tumor activity in vitro and in vivo^23^. Because IL-15 supports NK-cell metabolic fitness, we asked whether these cells could withstand hypoxic stress^29,33^.

We generated CD70-CAR-IL-15 NK-92 cells (**Fig. 2A**; CAR expression in Fig. **S2A**) and confirmed high CD70 surface expression on all three target cancer cell lines: Panc-1, HeLa, and Raji (**Fig. 2B**). CAR NK cells were pre-conditioned for 48 hours in either normoxia or hypoxia prior to co-culture. Hypoxia significantly reduced CD70-CAR-IL-15 NK cell-mediated killing across all targets (**Fig. 2C**). The magnitude of suppression was most pronounced in Raji and HeLa, while the effect on Panc-1 cells was more modest.

**Figure 2.**
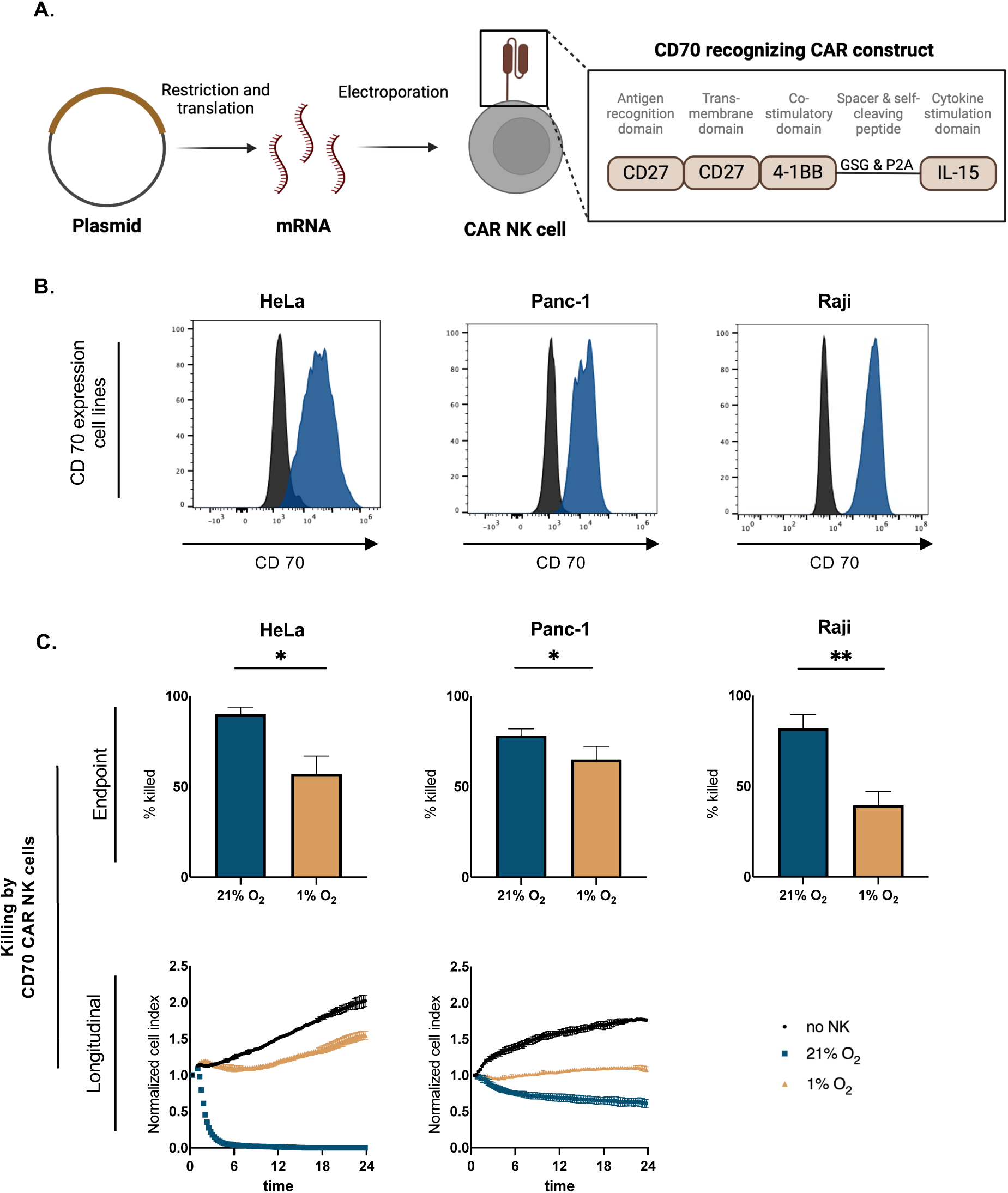
CD70 targeting CAR NK cells are rendered dysfunctional in hypoxia. A. Schematic representation of CAR mRNA synthesis and electroporation into NK cells, based on the protocol described by Van den Eynde *et al.*; B. CD70 surface expression on HeLa, Panc-1, and Raji target cell lines, as determined by flow cytometry, blue histograms represent CD70 staining and black histograms show fluorescence minus one (FMO) controls; C. Cytotoxicity of CD70-CAR-IL-15 NK cells against CD70^+^ targets under normoxic (21% O₂) and hypoxic (1% O₂) conditions. For HeLa and Panc-1, longitudinal killing was measured by xCELLigence (normalized cell index over 24 h), while Raji killing was assessed by flow cytometry after 4 h, bar charts show endpoint % target cell killing and curves show real-time cytolytic activity. *p < 0.05; **p < 0.01.

These results demonstrate that, even with IL-15 support, hypoxia imposes a suppressive effect on antigen-specific CAR NK cell function.

### Mdivi-1 treatment alleviates the adverse effects of hypoxia

As hypoxia impaired mitochondrial content and function, and IL-15-CAR engineering failed to restore cytotoxicity, we next investigated whether inhibiting the fission factor DRP1 could mitigate NK cell dysfunction. DRP1 has been implicated in hypoxia-induced mitochondrial fragmentation, contributing to impaired NK cell function in tumors^8^.

To test this, we used the pharmacological DRP1 inhibitor Mitochondrial Division Inhibitor 1 (Mdivi-1) **(Fig 3A)**. Mdivi-1 treatment restored K562 killing under hypoxia and prevented the loss of mitochondrial content (**Fig. 3B, C and Fig. S1B**). However, the specificity of this small molecule has been questioned, as it may affect cellular metabolic pathways beyond mitochondrial fragmentation^34^. Moreover, its clinical applicability remains limited due to the lack of safety data.

**Figure 3.**
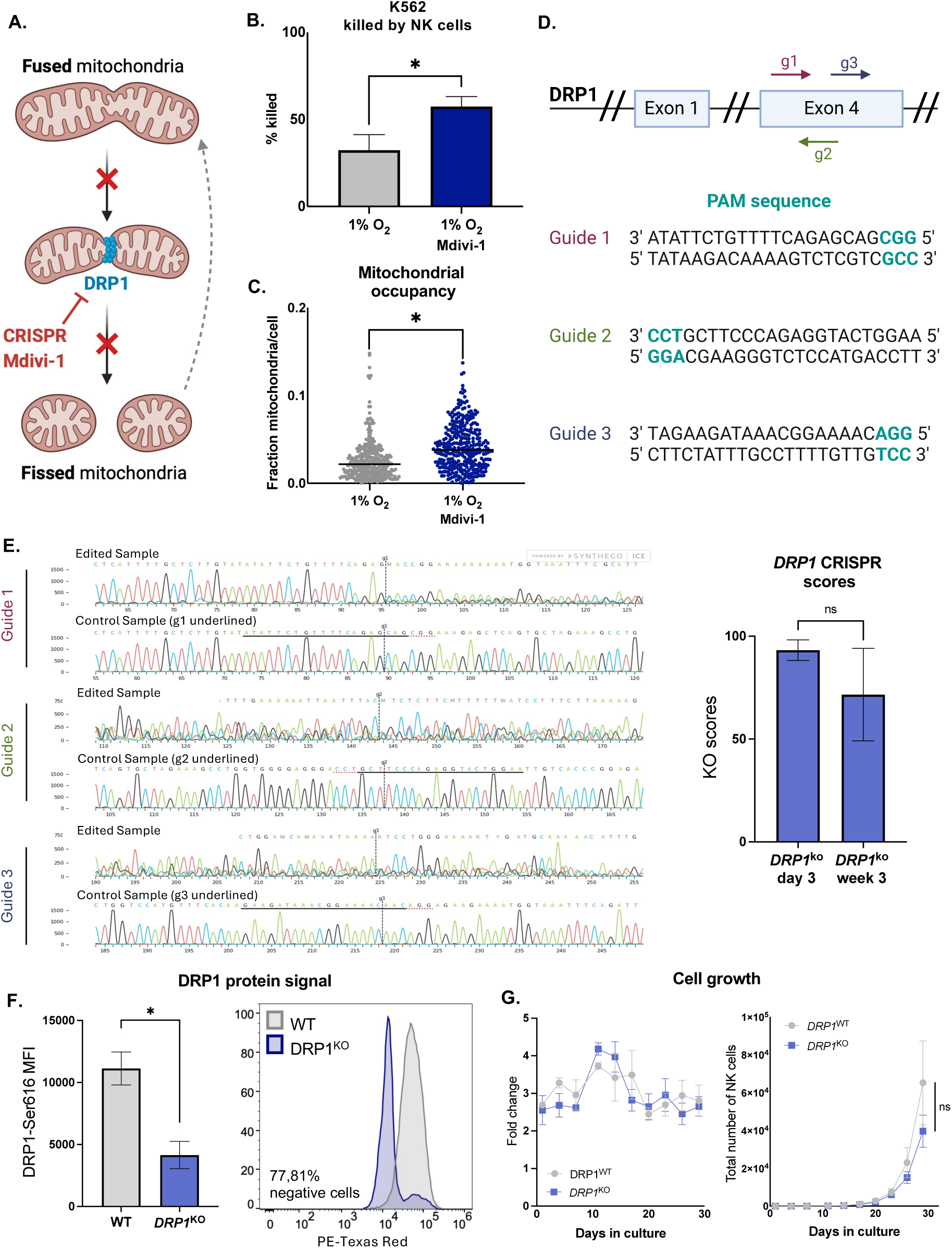
Mdivi-1 treatment and CRISPR-Cas9-mediated deletion of *DRP1* in NK cells. A Schematic representation of DRP1-mediated mitochondrial fission and its inhibition by Mdivi-1 or CRISPR-Cas9-mediated knockout (KO); B. Cytotoxic activity (% K562 cells killed) of NK cells cultured under hypoxia (1% O₂) with or without Mdivi-1 treatment; C. Mitochondrial occupancy in hypoxic NK cells with or without Mdivi-1, measured by confocal microscopy; D. Schematic of the *DRP1* gene with target sites for the three guide RNAs used for CRISPR editing; E. Representative Sanger sequencing chromatograms (left) for each guide RNA and corresponding KO egiciency scores (right) at day 3 and week 3 post-editing, analyzed with Synthego’s ICE tool; F. Flow cytometric quantification of DRP1 protein expression in DRP1^KO^ vs DRP1^WT^ NK cells.; G. Growth curves of DRP1^WT^ vs DRP1^KO^ NK cells. *p < 0.05; **p < 0.01; ns = not significant.

To overcome these limitations, we generated a stable KO of the *DRP1* gene using CRISPR-Cas9 technology. Using pre-formed Cas9 protein and a multi-guide RNA strategy targeting exon 4, we achieved high and durable KO efficiency, as confirmed by Sanger sequencing (**Fig. 3D, E**). Consequently, DRP1^KO^ NK cells showed reduced phospho-DRP1 levels. Importantly, DRP1-ablation maintained proliferation in NK cells at comparable rate to wild-type controls (**Fig. 3F, G**).

Hence, while DRP1 is implicated in hypoxia-induced mitochondrial and functional NK cell impairment, its loss does not compromise cell viability or proliferative capacity, setting the stage for functional assessment.

### DRP1^KO^ CAR NK cells retain their mitochondrial function and killing potential in hypoxia

DRP1^KO^ NK cells maintained mitochondrial integrity under hypoxic conditions, with mitochondrial occupancy preserved at levels comparable to normoxia (**Fig. 4A, Fig. S4A, B**). Mitochondrial membrane potential remained stable under hypoxia (**Fig. 4B**), while mtROS levels were still elevated in hypoxia (**Fig. 4C**).

**Figure 4.**
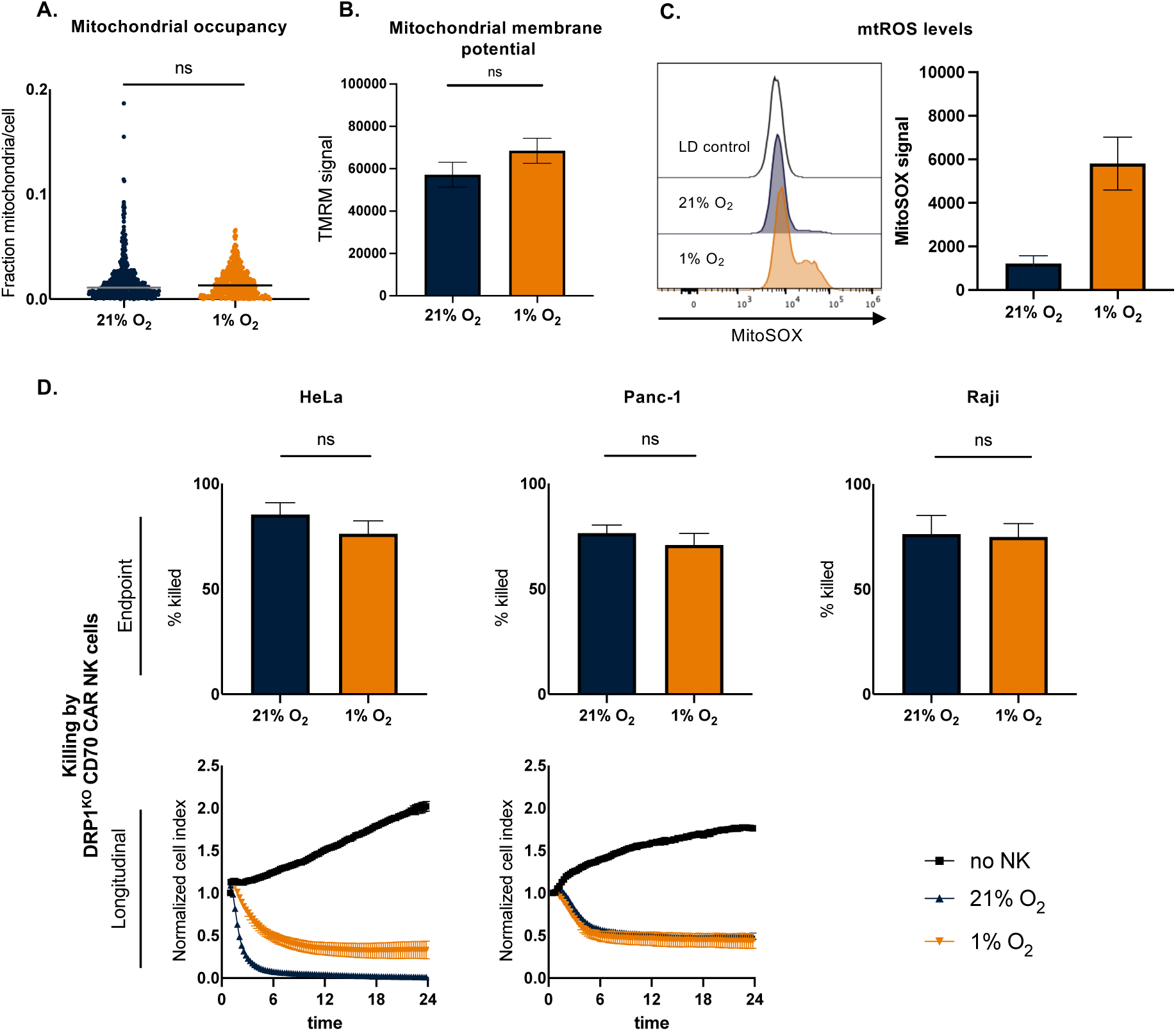
DRP1^KO^ NK cells retain mitochondrial integrity and cytotoxic function under hypoxia. A. Mitochondrial occupancy in DRP1^KO^ NK cells cultured in normoxic (21% O₂) or hypoxic (1% O₂) conditions, assessed by 3D confocal imaging, each point represents a single cell; B. Mitochondrial membrane potential measured by TMRM staining in DRP1^KO^ NK cells under normoxia or hypoxia, analyzed by flow cytometry; C. mtROS levels after normoxic/hypoxic culturing as determined by mitoSOX staining; D. Cytotoxic activity of DRP1^KO^ CD70-CAR-IL-15 NK cells against HeLa, Panc-1, and Raji target cells cultured in normoxic or hypoxic conditions, cytotoxicity was measured by xCELLigence (HeLa, Panc-1) and flow cytometry (Raji), upper panels show endpoint % killing and lower panels show longitudinal normalized cell index over 24 hours. ns = not significant.

To assess the functional relevance of this metabolic resilience, we engineered DRP1^KO^ NK cells to express the CD70-CAR-IL-15 construct (**Fig. S4C**) and evaluated their cytotoxic capacity under hypoxic conditions. DRP1^KO^ CAR NK cells maintained high cytotoxicity in both normoxic and hypoxic settings across all tested tumor cell lines (**Fig. 4D**).

In conclusion, DRP1^KO^ NK cells maintained mitochondrial content, membrane potential, and cytotoxic function under hypoxic conditions, supporting their potential to resist hypoxia-induced dysfunction.

### Despite DRP1 deletion, NK cells retain a general hypoxia-associated profile

Next, we investigated the transcriptional and metabolic profiles of DRP1^KO^ cells in hypoxic and normoxic conditions. The gene expression profile and top up- and downregulated Hallmark pathways were similar to the DRP1^WT^ cells in normoxic vs hypoxic conditions (**Fig. 5A, B**). However, a notable exception was the mTORC1 hallmark. While this pathway was downregulated in hypoxic DRP1^WT^ NK cells (**Fig. 1F**), it remained stable in DRP1^KO^ cells under the same conditions (**Fig. 5C**). This might be indicative of its preserved killing capacity.

**Figure 5.**
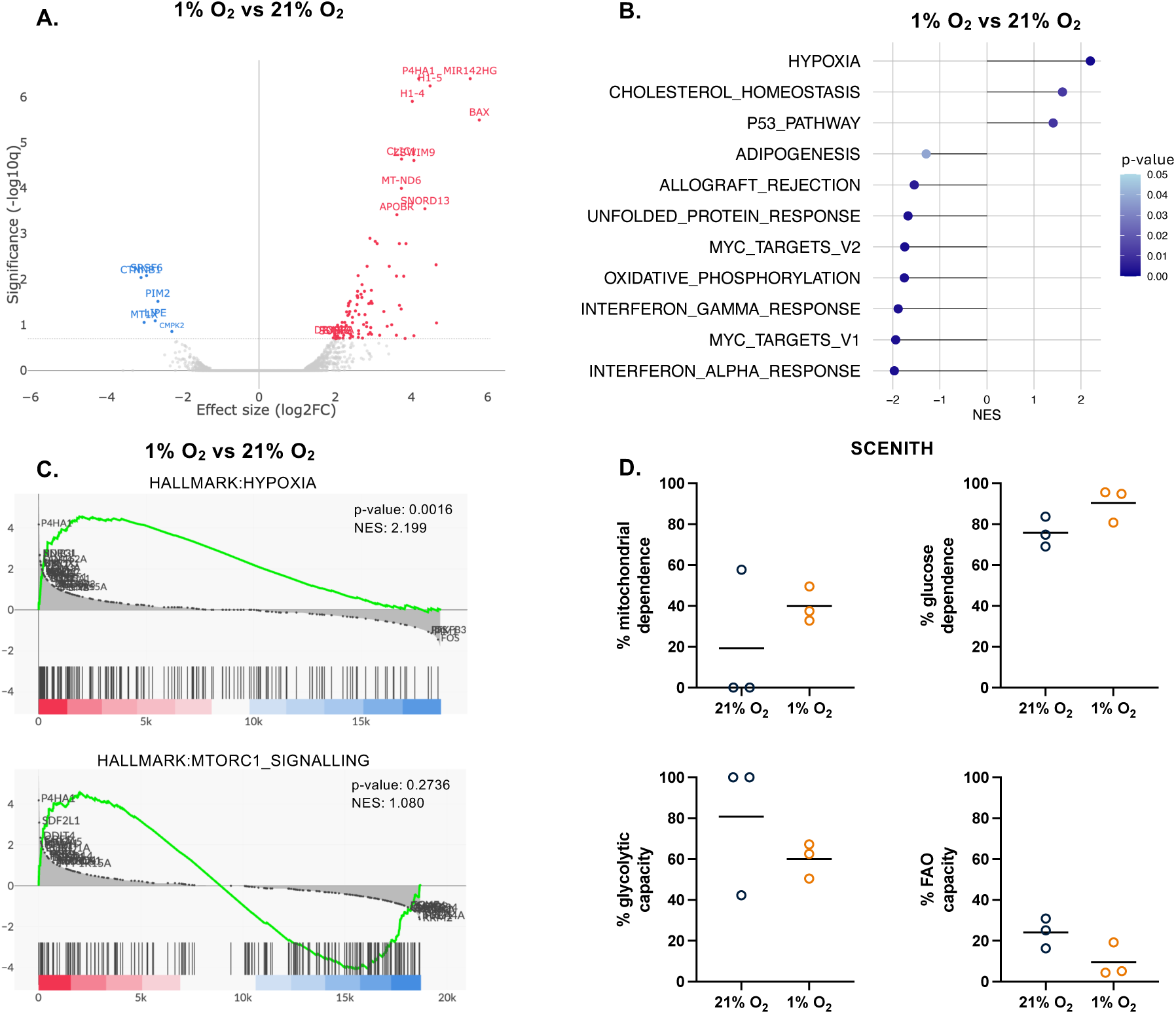
Transcriptomic and metabolic profiling of DRP1^KO^ NK cells under hypoxia. A. Volcano plot showing differentially expressed genes between DRP1^KO^ NK cells cultured in hypoxia (1% O₂) versus normoxia (21% O₂); significantly upregulated and downregulated genes are highlighted.; B. Hallmark gene set enrichment analysis displaying the top up- and downregulated pathways in hypoxic versus normoxic DRP1^KO^ NK cells.; C. Gene set enrichment analysis (GSEA) plots demonstrating increased expression of the hypoxia gene signature and no significant reduction in mTORC1 signaling in hypoxic DRP1^KO^ NK cells.; D. SCENITH-based analysis of metabolic activity in DRP1^KO^ NK cells under normoxia and hypoxia, showing corresponding metabolic phenotypes.

To determine whether DRP1 KO alters the metabolic response of NK cells to hypoxia, we performed SCENITH analysis on DRP1^KO^ cells cultured under normoxic and hypoxic conditions (**Fig. 5D**). Similar to wild-type cells, DRP1^KO^ NK cells displayed reduced translation in hypoxia (**Fig. S5**). Interestingly, despite the downregulation of OXPHOS-associated genes in the RNA-seq data, SCENITH revealed a trend toward increased mitochondrial dependence in hypoxia, consistent with the preserved mitochondrial membrane potential and mitochondrial content observed in DRP1^KO^ cells (**Fig. 5D**).

Overall, while hypoxia induces a hypoxic profile in DRP1^KO^ NK cells, a preserved mTORC1 signalling and increased mitochondrial dependence might enable the sustained killing capacity.

## Discussion

Hypoxia is a hallmark of solid tumors and is known to compromise the function of immune cells, including NK cells, by altering mitochondrial health and disrupting metabolic pathways^6,8,12^. Consistent with earlier findings, our results show that exposure to low oxygen levels leads to reduced mitochondrial content and membrane potential, elevated mtROS, and impaired NK cell cytotoxicity^8,12^. These functional impairments were also observed in NK cells engineered with a CD70-targeting CAR and IL-15. While CAR-based strategies and cytokine support enhance NK cell function in general, they appear insufficient to fully counteract hypoxia-induced dysfunction. Consistently, Kennedy *et al.* (2024) reported that IL-15 stimulation improved but did not fully preserve NK cell cytotoxicity under hypoxic conditions^12^. These observations underscore the need for strategies that more directly address the metabolic limitations imposed by the hypoxic tumor microenvironment.

Mitochondria are key regulators of cellular metabolism and immune cell fate^35^. They constantly undergo fission and fusion, reshaping their structure. Fission, mediated by DRP1 and FIS1, facilitates mitophagy, ROS production, apoptosis, and cell proliferation^36–41^. Fusion, controlled by MFN1, MFN2, and OPA1, forms tubular networks that enhance oxidative phosphorylation (OXPHOS), increase Ca²⁺ flux, and protect cells from stress^42–46^. In hypoxic regions of human tumors, tumor-infiltrating NK cells exhibit increased DRP1 activation, driven by sustained mTOR signaling^8^. This results in mitochondrial fragmentation, reduced mitochondrial polarization, increased mtROS, and impaired cytotoxicity, which is also observed in our study^8^.

Prompted by the essential role of mitochondria in NK cell biology and prior evidence implicating DRP1-mediated fission in NK cell dysfunction under hypoxia, we explored whether targeting DRP1 could protect NK cell function^8,47^. Our findings show that pharmacological DRP1 inhibition with Mdivi-1 restores mitochondrial defects and cytotoxicity. Consistent with our results, Zheng *et al.* (2019) showed that Mdivi-1 improved mitochondrial integrity and cytolytic activity of tumor-infiltrating NK cells exposed to hypoxia. However, due to concerns about the specificity and safety of Mdivi-1, we generated stable DRP1^KO^ NK cells using CRISPR-Cas9 technology. To our knowledge, this is the first report describing the successful generation and functional evaluation of DRP1^KO^ NK cells.

Previous work in neuronal cells showed that CRISPR-Cas9-mediated DRP1 KO protected against ferroptotic cell death by preserving mitochondrial morphology, membrane potential, and stabilizing redox balance^48^. In addition, pharmacological or genetic inhibition of DRP1 can exert anti-apoptotic effects, for example by blocking Bax-dependent cytochrome c release^49,50^. While we observed similarly preserved mitochondrial content and potential in DRP1^KO^ NK cells exposed to hypoxia, mtROS remained elevated and pro-apoptotic BAX expression was not fully suppressed, suggesting only partial protection against apoptotic signaling. In T cells, DRP1 deletion has been reported to skew cells toward a memory-like phenotype, at the expense of migration and cytoskeletal dynamics^51,52^. Whether similar trade-offs occur in NK cells remains unknown. The precise mechanisms by which DRP1^KO^ NK cells maintain cytotoxicity in hypoxia are not yet fully understood and may involve altered mitochondrial ultrastructure, fusion-fission balance, and changes in metabolic pathways.

Transcriptomic analysis showed that DRP1^KO^ and DRP1^WT^ NK cells had largely similar transcriptional processes under hypoxia, with one notable exception: mTORC1 signaling. While this pathway was downregulated in hypoxic DRP1^WT^ cells, it remained largely unchanged in DRP1^KO^ cells. Cytokine stimulation of murine and human NK cells results in robust mTOR activation, which is required for metabolic and functional responses, such as enhanced glycolysis and IFN-y production^29,53,54^. The precise mechanisms of mTOR regulation after IL-15 activation remain elusive in NK cells^55^. Given the role of mTORC1 in regulating NK cell metabolism and function, its preservation may contribute to the functional differences observed^29,54^. Additional work is needed to confirm this relationship. Interestingly, our findings contrast with those of Zheng *et al*., who reported heightened mTORC1 activation in NK cells under hypoxic conditions^8^. Several factors could account for this discrepancy, including differences in cell type, hypoxia duration, culture conditions, and the specific readouts used to assess mTORC1 activity. These contrasting observations highlight the complexity of mTORC1 regulation in immune cells and underscore the importance of context when interpreting pathway activity under hypoxic stress.

Recent studies have highlighted that ex vivo-expanded NK cells may naturally resist hypoxia-induced dysfunction. NK cells expanded using feeder-cell systems or antibody-based protocols have shown preserved cytotoxicity even after prolonged exposure to low oxygen conditions^12,56^. Feeder-expanded NK cells exhibited improved metabolic features, including a lower NAD⁺/NADH ratio, preserved AMP levels, and greater resilience to oxidative stress, contributing to enhanced metabolic stability^12^. A deeper understanding of the mitochondrial mechanisms and adaptations that confer this stability may offer opportunities to further optimize NK cell products, either by adopting expansion-based strategies or combining them with mitochondrial rewiring approaches like DRP1 modulation.

In conclusion, our findings show that hypoxia exerts wide-ranging effects on NK cell function, which cannot be fully rescued by cytokine support through CAR engineering alone. Given the central role of mitochondria in NK cell fitness, we focused on modulating mitochondrial dynamics. Our data suggest that disrupting DRP1 in NK cells can help preserve mitochondrial function and cytotoxicity in hypoxia. Such mitochondrial modulation might be required for effective functioning of CAR NK cells in hypoxic solid tumors.

## Acknowledgements

This work was supported by Research Foundation Flanders (G040120N; J.M. 11PHM24N) and by the University of Antwerp (BOF and IOF grant numbers FFB230049, FFB230316, FFI230209). T.V. received support from Kom op tegen Kanker (Stand up to Cancer), the Flemish cancer society (projectID: 13911). P.V. received support from the Alzheimer Research Foundation (SAO-FRA) (project 2023004). J.D.W. received support from the ME TO YOU foundation (grant number OZ8546). Part of the research was supported by donations from different donors, including Willy Floren, Dedert Schilde vzw and the Vereycken family.

---

**Figure S1.**
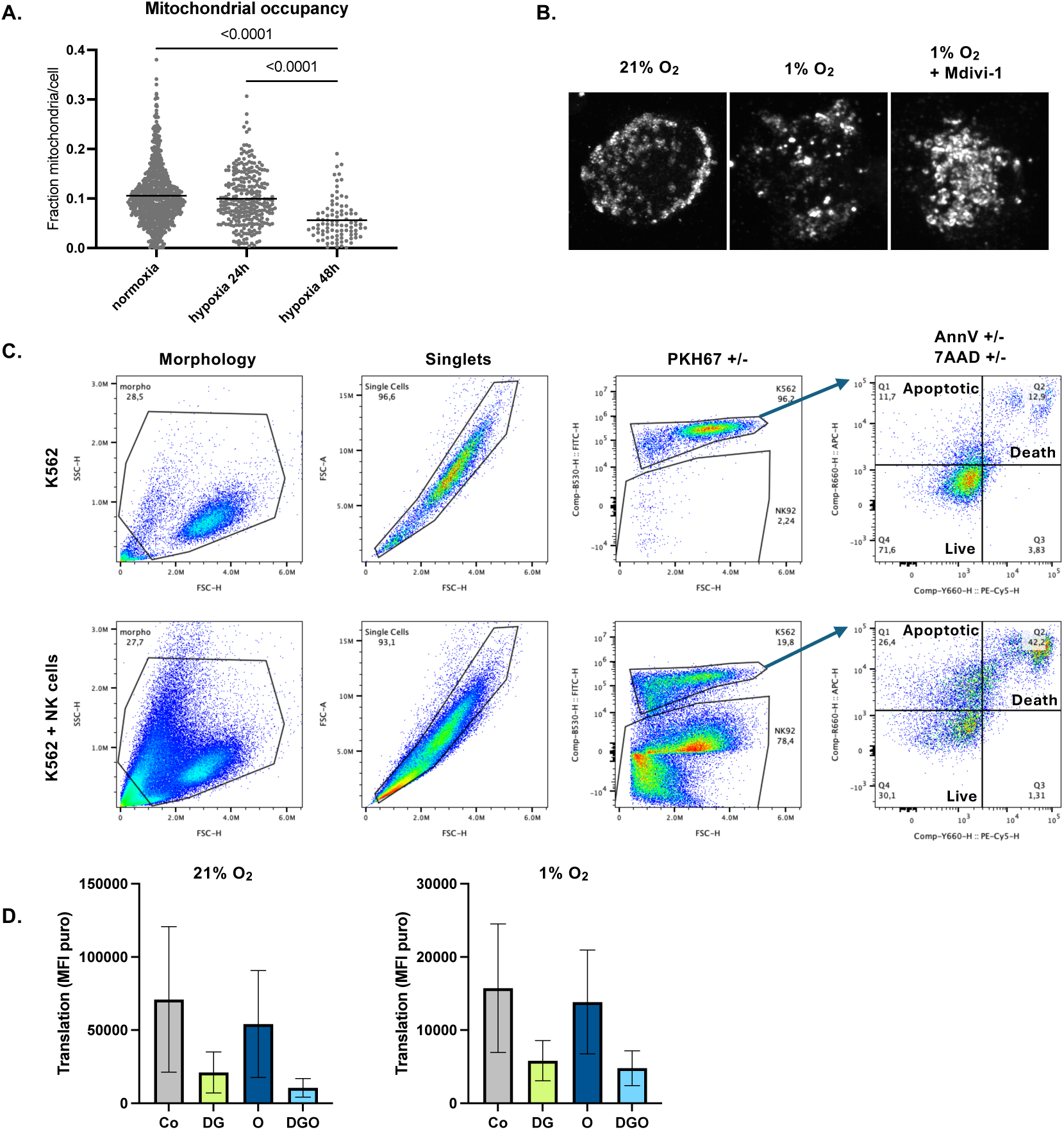
Features of NK cells in normoxia and hypoxia and gating strategy. A. Mitochondrial occupancy over time in hypoxic culturing, determined by confocal microscopy; B. Representative images for confocal microscopy quantifications; C. Gating strategy for co-culture assays with K562 cells and NK cells on flow cytometry; D. Assessment of protein translation in NK cells cultured in normoxia or hypoxia for 48 hours, as measured by puromycin incorporation (SCENITH assay). Cells were treated with control (Co), 2-deoxyglucose (DG), oligomycin (O), or the combination (DGO).

**Figure S2.**
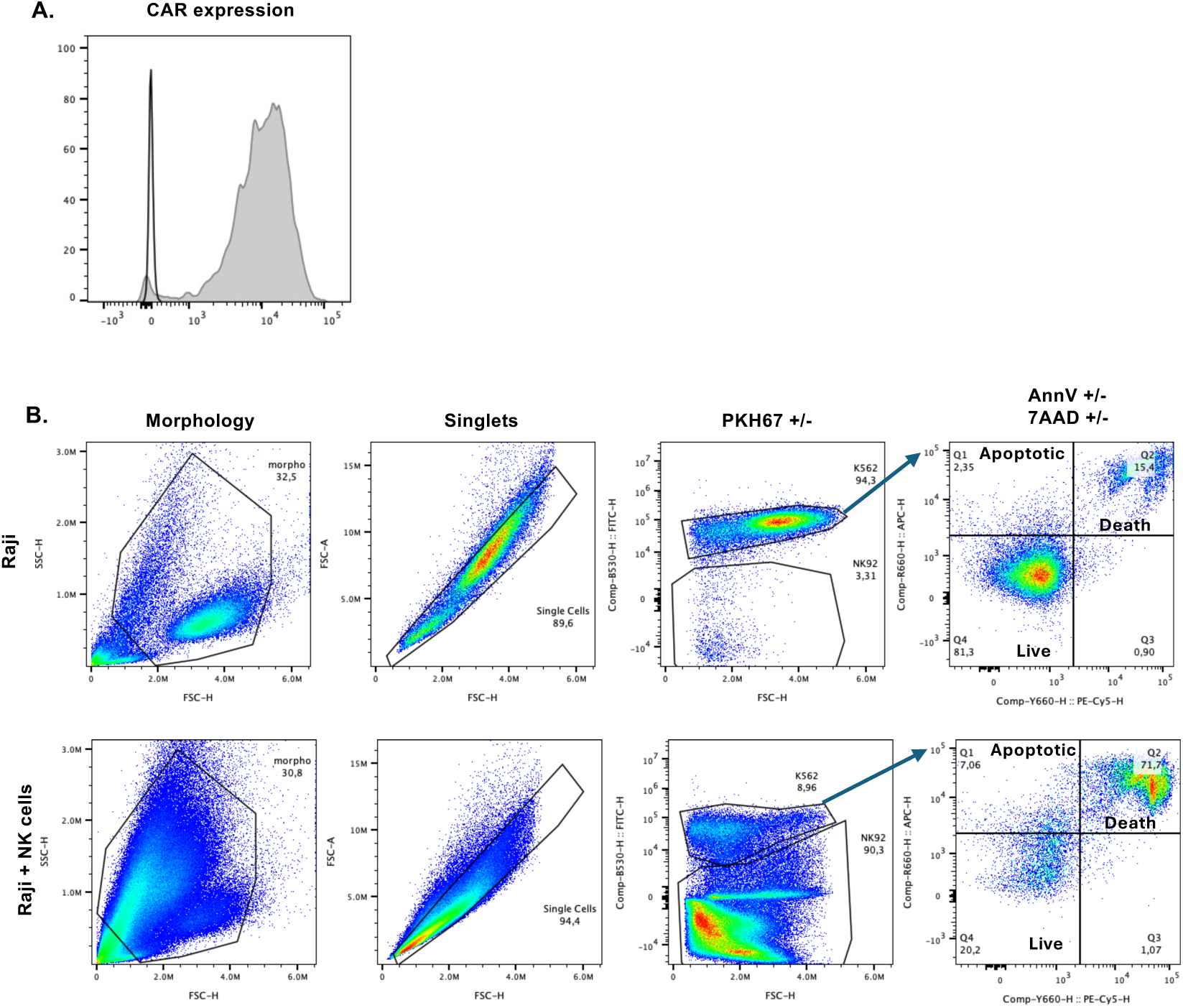
Features of CAR NK cells and gating strategy. A. Representative histograms of CAR expression on CAR NK cells determined by flow cytometry; B. Gating strategy for co-culture assays with Raji cells and CD70-CAR-IL15 NK cells on flow cytometry.

**Figure S3.**
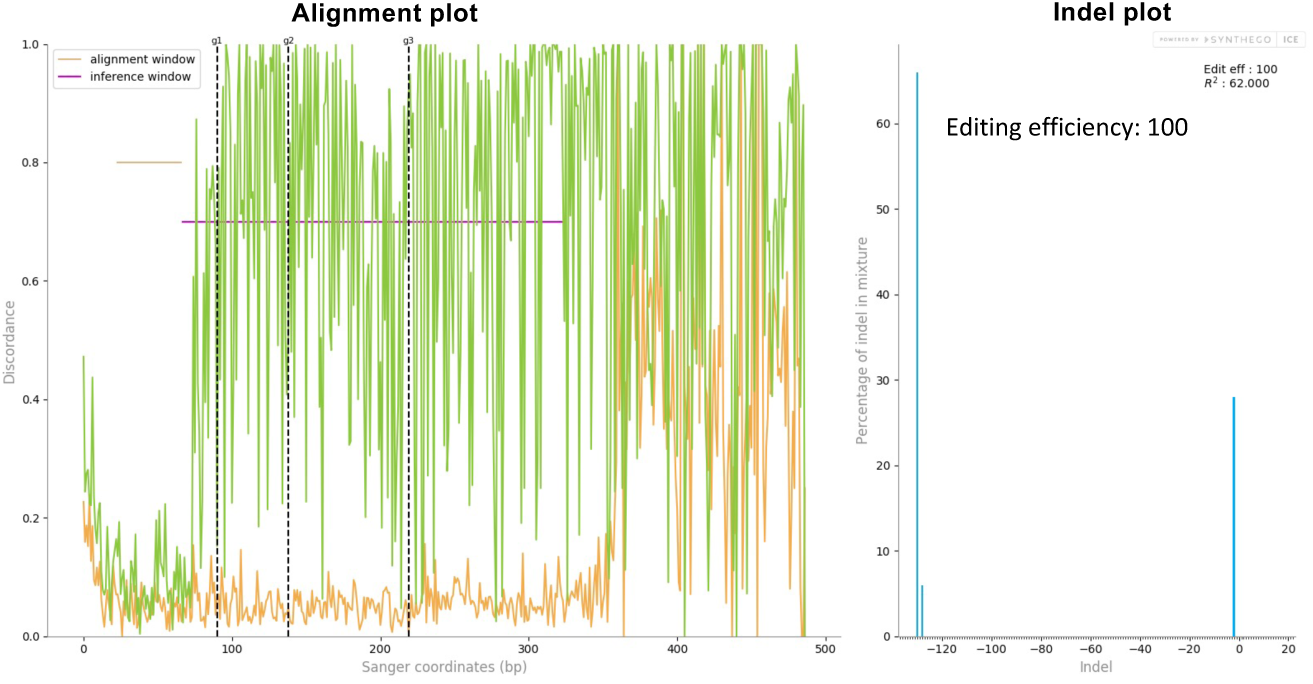
Analysis of Editing ERiciency of the DRP1 gene in NK cells. Determined by Sanger sequencing and ICE analysis. (left) An alignment plot displaying the aligned control (orange) and edited (green) sequences; (right) an indel plot depicting the anticipated range of insertions and deletions within the edited gene locus and the editing egiciency.

**Figure S4.**
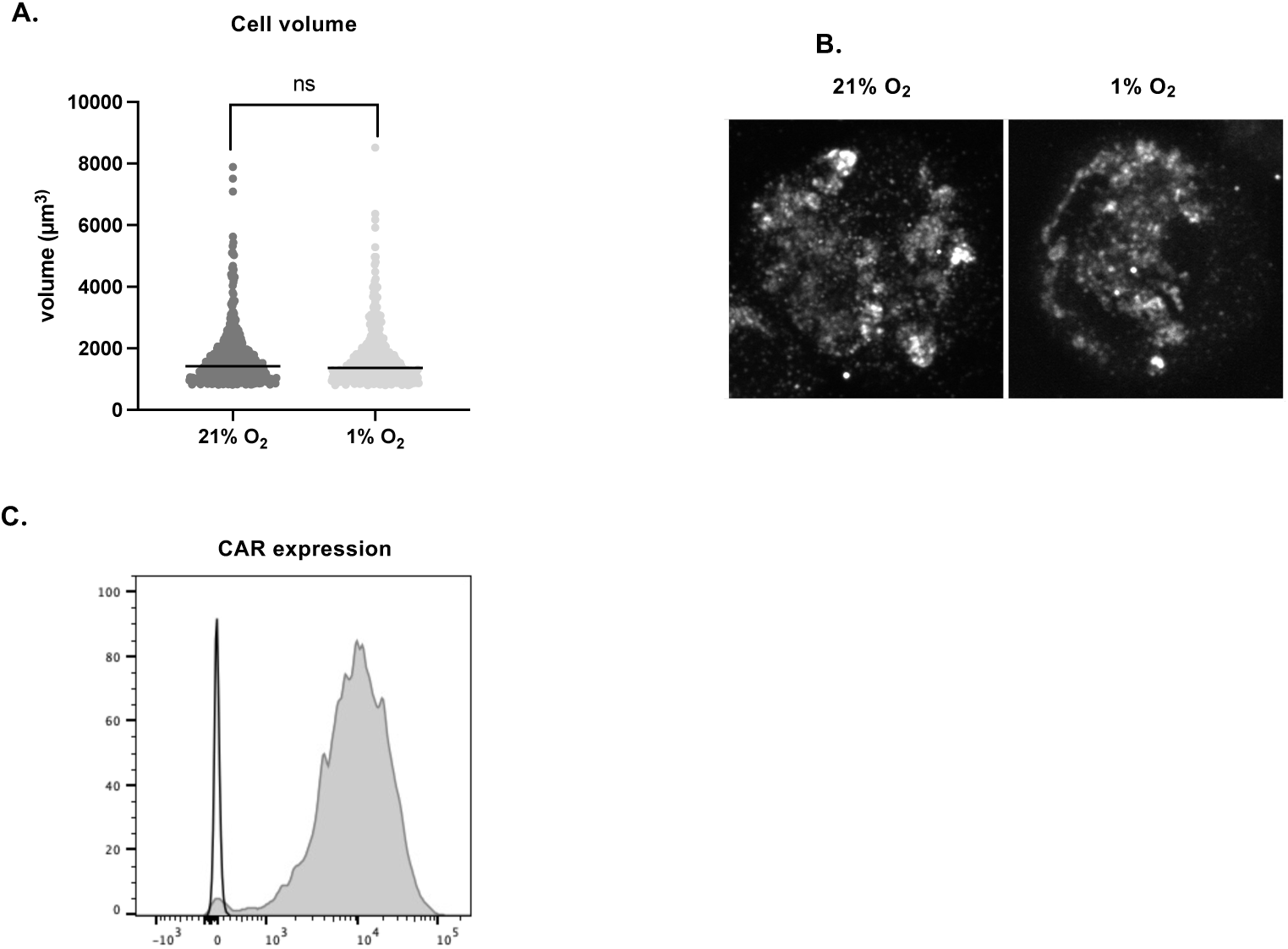
**Features of DRP1^KO^ NK cells**. A. Cell volume determined by confocal microscopy; B. Representative images used for quantification of confocal microscopy; C. Representative histograms of CAR expression on DRP1^KO^ CAR NK cells determined by flow cytometry.

**Figure S5.**
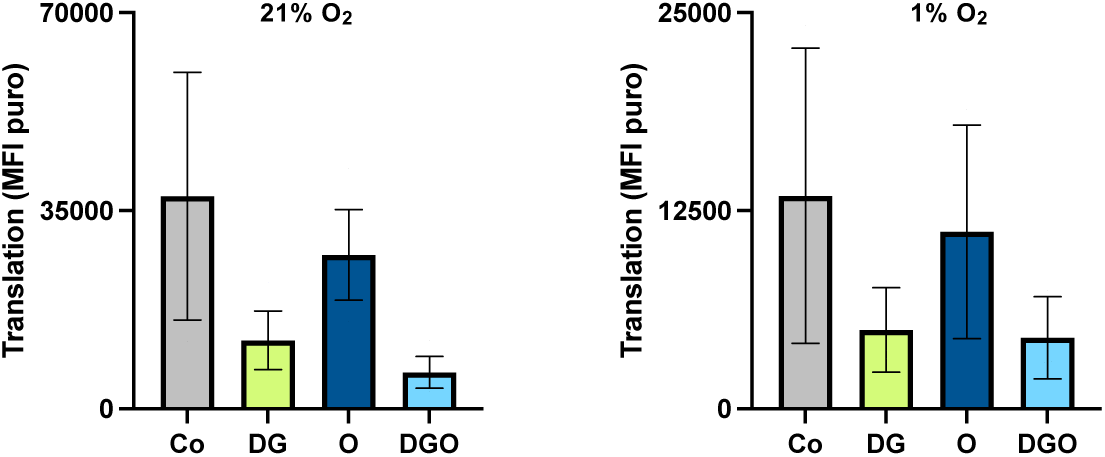
Translation in DRP1^KO^ cells. Assessment of protein translation in NK cells cultured in normoxia or hypoxia for 48 hours, as measured by puromycin incorporation (SCENITH assay). Cells were treated with control (Co), 2-deoxyglucose (DG), oligomycin (O), or the combination (DGO).

